# Lipidomic analysis of roseobacters of the pelagic RCA cluster and their response to phosphorus limitation

**DOI:** 10.1101/2020.11.21.392431

**Authors:** Eleonora Silvano, Mingyu Yang, Mathias Wolterink, Helge-Ansgar Giebel, Meinhard Simon, David J Scanlan, Yanlin Zhao, Yin Chen

**Author notes:** Correspondence to Dr Y Chen).

## Abstract

The marine roseobacter-clade Affiliated cluster (RCA) represents one of the most abundant groups of bacterioplankton in the global oceans, particularly in temperate and sub-polar regions. They play a key role in the biogeochemical cycling of various elements and are important players in oceanic climate-active trace gas metabolism. In contrast to copiotrophic roseobacter counterparts such as *Ruegeria pomeroyi* DSS-3 and *Phaeobacter* sp. MED193, RCA bacteria are truly pelagic and have smaller genomes. We have previously shown that RCA bacteria do not appear to encode the PlcP-mediated lipid remodelling pathway, whereby marine heterotrophic bacteria remodel their membrane lipid composition in response to phosphorus (P) stress by substituting membrane glycerophospholipids with alternative glycolipids or betaine lipids. In this study, we report lipidomic analysis of six RCA isolates. In addition to the commonly found glycerophospholipids such as phosphatidylglycerol and phosphatidylethanolamine, RCA bacteria synthesise a relatively uncommon phospholipid, acylphosphatidylglycerol, which is not found in copiotrophic roseobacters. Instead, like the abundant SAR11 clade, RCA bacteria upregulate ornithine lipid biosynthesis in response to P stress, suggesting a key role of this aminolipid in the adaptation of marine heterotrophs to oceanic nutrient limitation.

## Introduction

The marine roseobacter group of *Alphaproteobacteria* comprises an ecologically important group of marine bacteria involved in the biogeochemical cycling of carbon, nitrogen and sulphur (Buchan et al., 2014; Luo & Moran, 2014). Roseobacters are metabolically diverse, being able to use an array of organic molecules, perform anoxygenic photosynthesis and produce secondary metabolites (Buchan et al., 2014; Brinkhoff et al., 2008). Arguably however, they are most well known for their role in marine trace gas formation, including the metabolism of methylated sulfur compounds like dimethylsulfide and dimethylsulfoniopropionate, and methylated amines, which has global significance (Lidbury et al., 2014; Mausz & Chen, 2019; Curson et al., 2011).

Using model marine roseobacters, we have previously studied membrane lipids in *Ruegeria pomeroyi* DSS-3 and *Phaeobacter* sp. MED193 (Sebastian et al., 2016; Smith et al., 2019). In addition to glycerophospholipids such as phosphatidylglycerol (PG) and phosphatidylethanolamine (PE), these marine roseobacters have several lipids which are not widely reported, including amino-acid containing lipids like glutamine lipid (Smith et al., 2019). Many of these roseobacters are capable of remodelling their membrane lipid composition in response to environmental change, such as adaptation to P stress through a phospholipase (PlcP)-mediated lipid renovation pathway by substituting glycerophospholipids with non-phosphorus containing surrogate lipids (Sebastian et al., 2016). Indeed, some roseobacter clade bacteria, including *Phaeobacter* sp. MED193 can produce betaine-containing membrane lipids to replace glycerophospholipids in response to phosphorus limitation (Sebastian et al., 2016).

In contrast to the aforementioned strains, some marine roseobacters are truly pelagic with small genomes, which are known to be numerically abundant in marine surface waters, along with several other bacterial groups, notably the SAR11 clade (Luo & Moran 2014; Luo & Moran 2015; Sun et al., 2017; Giebel et al., 2011; Giovannoni 2017).

One of such cosmopolitan pelagic roseobacter groups is the RCA (roseobacter clade affiliated cluster) group which is particularly prevalent in the bacterioplankton in temperate and polar regions of the oceans, where they can reach up to 35% of total bacterial counts (Selje et al., 2004; Giebel et al., 2009; 2011; Billerbeck et al., 2016; Zhang et al., 2016). RCA group bacteria were first isolated from the North Sea represented by the type strain *Planktomarina temperata* RCA23 (Giebel et al., 2013). Other RCA isolates include *Roseobacter* sp. LE17, isolated from an algal culture off the coast of California (Mayali et al., 2008), three strains (IMCC1909, IMCC1923, IMCC1933) isolated from the Yellow Sea (Giebel et al., 2013) and, more recently, three RCA strains obtained using high-throughput dilution-to-extinction culturing method using seawater from the East China Sea (strains FZCC0023, FZCC0040 and FZCC0043) (Zhang et al., 2019). The genome sequence of strains RCA23 and LE17 has been reported and, interestingly, neither genome contains the genes involved in PlcP-mediated lipid remodelling, which are prevalent in other marine roseobacters (Sebastian et al., 2016). Furthermore, three single-cell amplified genomes (SAGs) belonging to the RCA were recently reported, none of which encodes the PlcP enzyme (Sun et al., 2017). This prompted us to characterise the intact membrane lipids in this ecologically important group of pelagic marine bacteria.

## Results and Discussion

We brought together six RCA strains isolated from geographically distinct locations around the world (**Figure 1**). Due to their unique growth requirements it was not feasible to use the same growth medium to cultivate all six strains. Indeed, genome sequence and comparative genomics studies have previously shown that each RCA strain has a unique requirement for vitamins (Voget et al., 2015; Zhang et al., 2019). Thus, strains RCA23, LE17 and IMCC1933 were cultivated in a diluted marine broth medium (40% w/v marine broth in seawater), whereas the strains isolated from the East China Sea (strains FZCC0023, FZCC0040 and FZCC0043) do not grow in marine broth or diluted marine broth medium. Instead, they were grown in autoclaved natural seawater amended with mixed vitamins, ammonium (1 mM), phosphate (100 μM), iron and a mixed carbon source (Zhang et al., 2019). To determine the impact of P depletion on the lipid composition in these latter three RCA strains, they were also grown on the same natural seawater amended medium without adding additional phosphate. The final cell density of these six RCA strains before harvest is shown in **Figure 1B**. Without phosphate, the final cell density only reached ~10^7^ cells/ml whereas with phosphate amended, they reached ~10^8^ cells/ml or more (**Figure 1B**), suggesting P can be a limiting factor for their growth.

**Figure 1.**
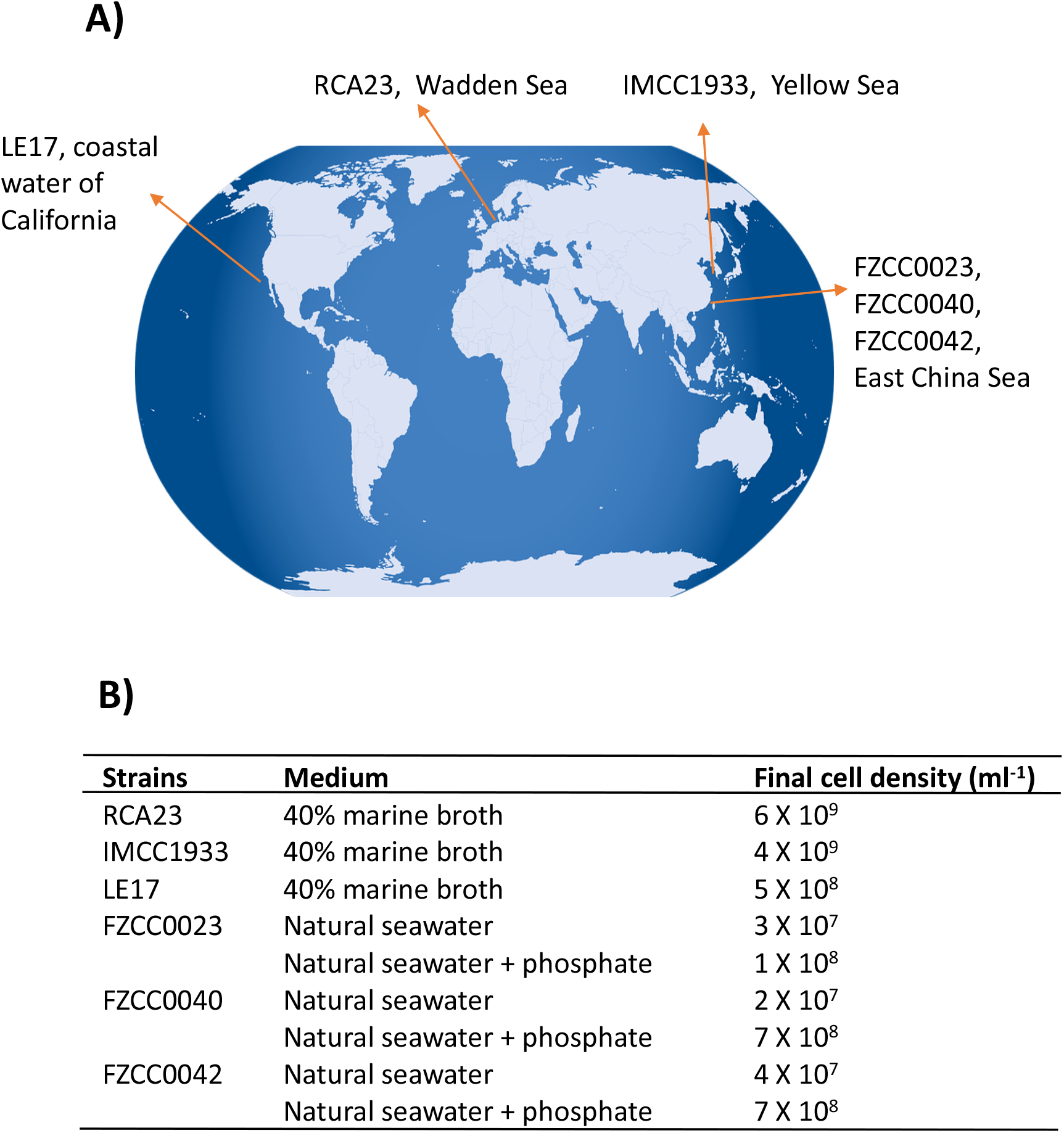
RCA isolates used in this study **A)** Geographical location of the six RCA strains; **B)** Culture medium used for each strain and the final cell density before harvesting.

The intact membrane lipids were extracted from these six bacterial cultures using a modified Folch method and their lipid composition analysed by liquid chromatography coupled with mass spectrometry (LC-MS) in both positive (+ve) and negative (-ve) ionisation mode (Smith et al., 2019). These lipids were separated using a hydrophilic interaction column prior to MS fragmentation and subsequent identification and quantification. Overall, four major phospholipids (phosphatidic acid (PA), phosphatidylglycerol (PG), phosphatidylethanolamine (PE), and acyl-PG (APG)) and one aminolipid (ornithine-containing) were consistently identified in these RCA bacterial isolates (**Table 1**). A representative LC-MS chromatograph of lipids from the RCA type strain (RCA23) is presented in **Figure 2**. The major fatty acids in these glycerophospholipids are C16:1, C18:1 and C19:1 (Table S1) although their composition varies depending on the individual lipids (top panel). This is consistent with previous estimations of fatty acid methyl esters by gas chromatography, showing that the two monounsaturated C16:1 and C18:1 species accounted for more than 70% of the total fatty acids in strain RCA23 (Giebel et al., 2013).

**Table 1.** Major lipids identified from RCA bacteria

| Retention time | Lipid class | m/z | Identity | Fatty acid species |
| --- | --- | --- | --- | --- |
| 3.6 min | Acyl-PG (APG) | 925.8 | [M-H]<sup>-</sup> | C18:1/C16:1/C12:1 |
|  |  | 953.8 | [M-H]<sup>-</sup> | C18:1/C18:1/C12:1 |
|  |  | 967.8 | [M-H]<sup>-</sup> | C18:1/C19:1/C12:1 |
| 6.5 min | PG | 745.7 | [M-H]<sup>-</sup> | C16:1/C18:1 |
|  |  | 773.7 | [M-H]<sup>-</sup> | C18:1/C18:1 |
|  |  | 787.7 | [M-H]<sup>-</sup> | C19:1/C18:1 |
| 11.3 min | PA | 729.7 | [M+Acetate]<sup>-</sup> | C16:1/C18:1 |
|  |  | 757.7 | [M+Acetate]<sup>-</sup> | C18:1/C18:1 |
|  |  | 771.7 | [M+Acetate]<sup>-</sup> | C19:1/C18:1 |
| 11.8 min | PE | 714.7 | [M-H]<sup>-</sup> | C16:1/C18:1 |
|  |  | 742.7 | [M-H]<sup>-</sup> | C18:1/C18:1 |
|  |  | 756.7 | [M-H]<sup>-</sup> | C19:1/C18:1 |
| 13.0 min | OL | 703.7 | [M-H]<sup>-</sup> | OL3-OH20:1/C18:1 |
|  |  | 717.7 | [M-H]<sup>-</sup> | OL3-OH20:1/C19:1 |

PG: phosphatidylglycerol; APG, acyl-phosphatidylglycerol; PA: phosphatidic acid; PE: phosphatidylethanolamine; OL, ornithine lipid.

**Figure 2.**
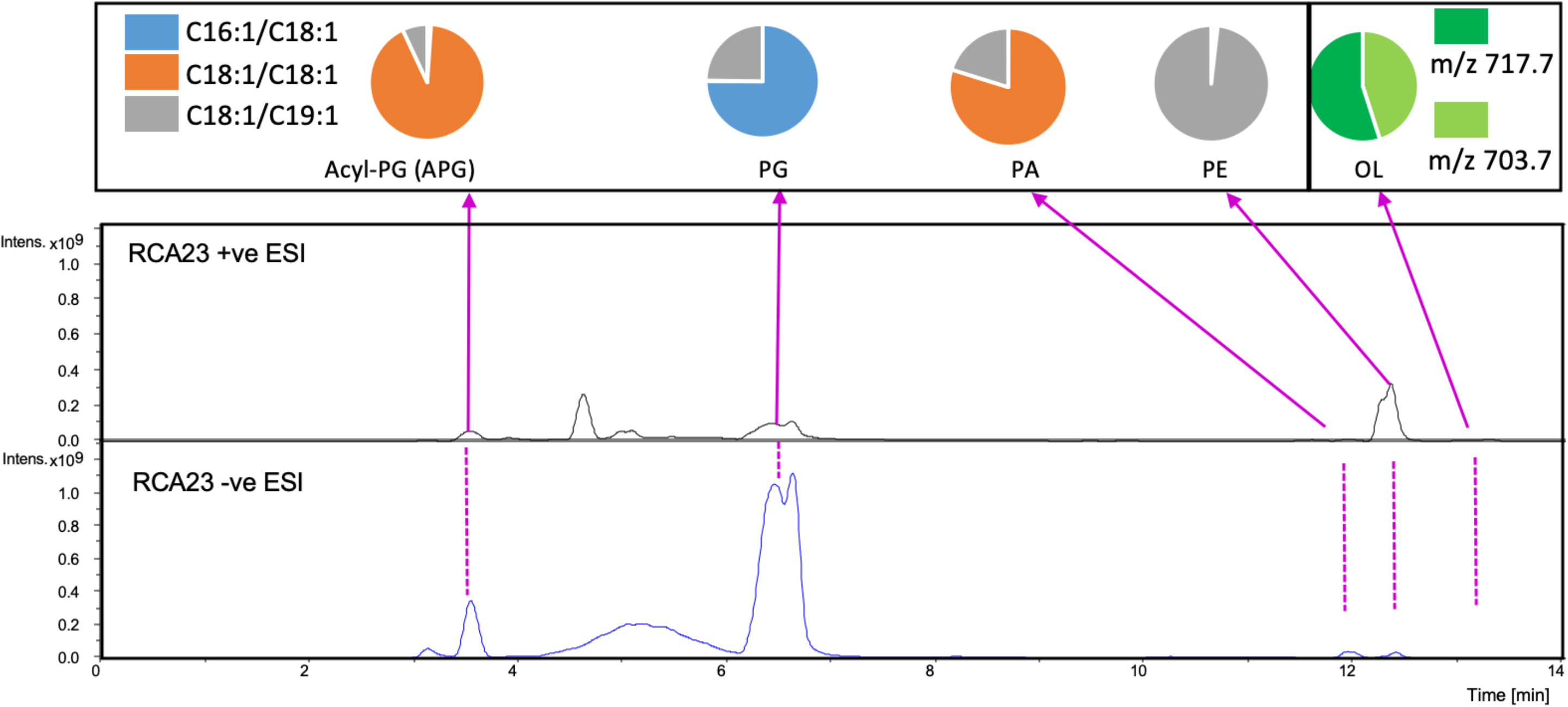
Major lipids detected in the RCA bacterium, type strain RCA23. An overview of the LC-MS chromatogram showing the major lipids extracted from strain RCA23. Lipids were analysed in both negative (- ve) and positive (+ ve) ionisation mode. The pie chart above each lipid class shows the major intact lipid species within the class. The colours represent the sn-1/sn-2 fatty acid composition in the phospholipids. Two major ornithine lipids were found in negative ionisation mode (see Table 1) with a mass-to-charge ratio of 703.7 and 717.7, respectively. A list of all lipids detected in other RCA strains are shown in supplementary Table S1. PA, phosphatidic acid; PG, phosphatidylglycerol; PE, phosphatidylethanolamine; OL, ornithine lipids; APG, acyl-PG.

In addition to PG, PE and a small amount of PA, we also observed phospholipids that eluted at 3.9 mins with a m/z ranging from 925.8-967.8 (**Table 1, Figure 2**). To elucidate the identity of these lipids, we isolated the major ion species with a m/z of 925.8 for further fragmentation analysis in the negative (-ve) ionisation model (**Figure 3**). The MS^n^ fragmentation pattern suggests that this lipid is a modified C34:2 PG (C16:1/C18:1) by acylation (APG) with a third acyl fatty acid R3’ of C12:1. Interestingly, 12-carbon fatty acids were previously found in RCA23 using gas chromatography analysis of fatty acid methyl esters (Giebel et al., 2013). APG lipids are rarely reported as a major lipid in bacteria (Yague et al., 1997; Hines & Xu 2019; Luo et al., 2018). Indeed, APG lipids were not detected in our previous studies of the lipidomes of marine roseobacters, including *Ruegeria pomeroyi* DSS-3 and *Phaeobacter* sp. MED193 (Sebastian et al., 2016; Smith et al., 2019) nor in pelagic SAR11 clade bacteria (Carini et al., 2015). APG is produced as a minor lipid in *E. coli* through direct acylation of the headgroup of PG using an acyl donor, with the outer membrane lipase PldB involved in generating this donor (Nishijima et al., 1975; Hines & Liu, 2019). Indeed, the genome sequences of RCA isolates RCA23 and LE17 also contain a PldB homologue showing 30% sequence identity (e^-20^) to that of *E. coli,* suggesting that APG in RCA strains is synthesized in a similar manner. However, it should be noted that the role of PldB is not limited to APG biosynthesis. Indeed, PldB of *Sinorhizobium meliloti* (SMc04041) also has thioesterase activity which is able to hydrolyse palmitoyl-CoA (Sahonero-Canavesi et al., 2015). Clearly the pathway for APG biosynthesis in RCA bacteria warrants further investigation.

**Figure 3.**
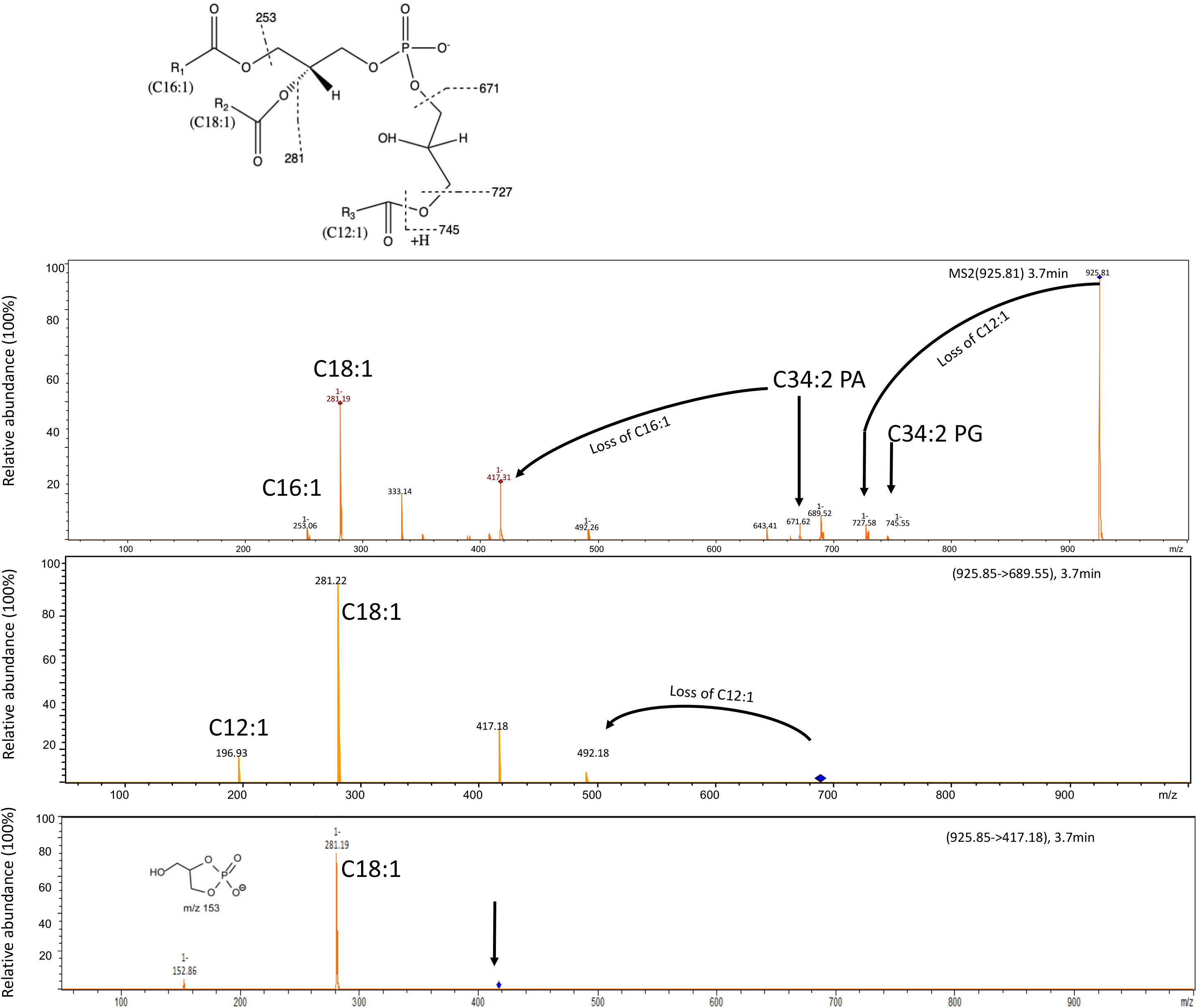
Fragmentation pattern of APG lipids eluted at 3.6 min and the proposed structure of the major APG lipids in these RCA bacterial isolates. Three APG lipids (Table 1) are found in all RCA strains with a mass-to-charge ratio of 925.8, 953.8 and 967.8, respectively. The mass spectra shown represents the APG lipids from the type strain RCA23. The dominant APG lipid species with a m/z ratio of 925.8 was selected for further fragmentation (MS^2^) which produces a lyso-PA species (m/z 417). This lyso-PA was further fragmented (MS^3^) to produce a C18:1 fatty acid and a characteristic ion of m/z 153. The mass difference between APG and the corresponding PG was 180.3, suggesting that the R3’ chain is C12:1 fatty acid.

In addition to these phospholipids, these RCA isolates also produced aminolipids. Thus, ornithine lipid (OL) eluted at 13 min, a feature consistent with the production of OLs in *R. pomeroyi* DSS-3 and *Phaeobacter* sp. MED193 (Smith et al., 2019). OLs in these RCA isolates comprise two major species of m/z 703.5 and 717.5 respectively, and their fragmentation pattern is consistent with a fatty acid composition of 3-hydroxyl20:1/C18:1 and 3-hydroxyl20:1/C19:1, respectively. Like *R. pomeroyi* DSS-3, the genomes of RCA isolates RCA23 and LE17, and SAGs AB-661-I11 and AB-661-L17, all contain a two gene cluster *olsA* and *olsB* encoding the *O*-acetyltransferase and *N*-acetyltransferase, respectively, which is likely responsible for the synthesis of OL. Glutamine lipid, which was observed in both *R. pomeroyi* DSS-3 and *Phaeobacter* sp. MED193, was not found in these RCA bacteria despite the fact that the *glsB* gene, which was previously shown to be essential for glutamine lipid biosynthesis in *R. pomeroyi* DSS-3, is present in their genomes (Smith et al., 2019).

To gain an insight into the role of P availability in modifying the lipidome of RCA bacteria, we used the three strains isolated from the East China Sea (strains FZCC0023, FZCC0040 and FZCC0043) and grew them in autoclaved natural seawater amended with 100 μM phosphate or without phosphate, as described previously (see Zhang et al., 2019). Analysis of the lipidome revealed no statistical difference (student *t*-test, *p*=0.31) in the relative abundance of the APG lipids. However, OLs were significantly more abundant (student *t*-test, *p*=0.01) in the P-stressed cultures, suggesting a role of OLs in response to P stress (**Figure 4**). It should be noted that, due to the lack of commercial standards of OL, only the relative abundance of OL/PG is analysed in this study, which does not take into account the potential differences in ionisation efficiency amongst different lipid classes. Nevertheless, a similar response of up-regulation of OLs in response to P stress was also observed in the marine heterotroph SAR11 strain HTCC7211 (Carini et al., 2015; Sebastian et al., 2016). In SAR11 clade bacteria, lipid renovation in response to P stress also involves the formation of several glycolipids, which are synthesized to replace glycerophospholipids. A bifunctional glycosyltransferase (Agt) is responsible for the formation of monoglycosyldiacylglycerol (MGDG) and glucuronic acid diacylglycerol (GADG). However, no homologues of Agt were present in the genomes of RCA clade bacteria and these glycolipids were also absent in their lipidomes in our analysis.

**Figure 4.**
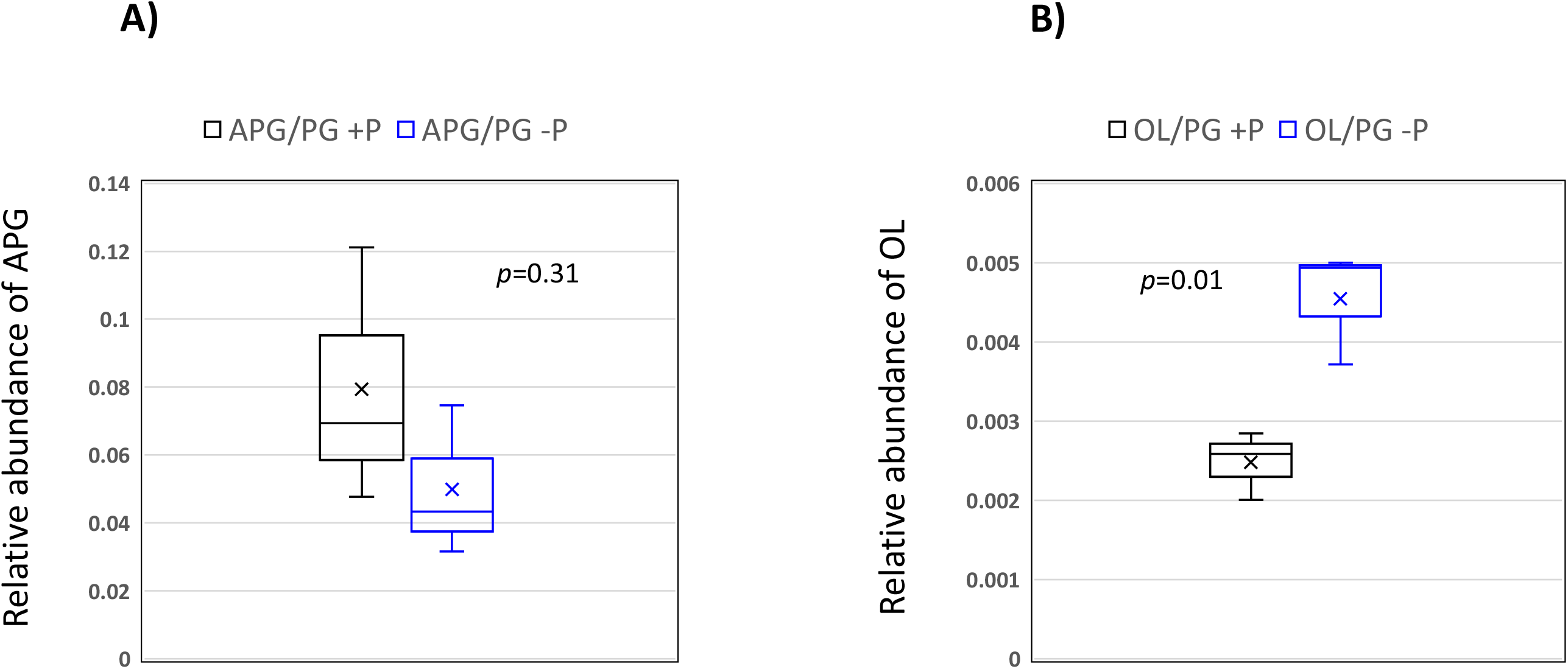
Box-whisker plot of the relative abundance of (A) APG and (B) OL lipids (relative to PG) in medium with and without phosphorus supplementation. The line in the middle of the box represents the mean whereas the symbol x represents the median. Data were from the three isolates (n=3) obtained from the East China Sea (strains FZCC0023, FZCC0040 and FZCC0043). Student-*t* tests were performed in Excel. PG, phosphatidylglycerol; OL, ornithine lipids; APG, acyl-PG.

Together, the data presented in this study reports the lipidome of six strains of RCA bacteria isolated from widely different geographical locations. These bacteria produce a relatively uncommon phospholipid, APG, although this lipid does not appear to play a role in the response to P stress. On the other hand, all RCA bacteria produce an ornithine-containing aminolipid which was significantly up-regulated under P stress. How these OLs facilitate membrane homeostasis in response to abiotic and biotic stresses certainly warrants further investigation.

## Materials and Methods

### Bacterial strains and cultivation

Strains RCA23 (North Sea), IMCC1933 (Yellow Sea) and LE17 (Pacific, West coast) were grown in 50 ml 40% (w/v) marine broth medium at 15°C in light-dark cycles for 20 days as described by Giebel et al. (2013). End point cell density was obtained by using a BD Accuri C6 flow cytometer (BD Biosciences, San Jose, CA, USA) after the protocol of Giebel et al. (2019). Cell pellets were harvested from a 50 ml culture by centrifugation at 4°C. Cells were stored at −80°C prior to lipid extraction.

Strains FZCC0023, FZCC0040 and FZCC0043 were isolated from the East China Sea and do not grow in marine broth or diluted marine broth medium. The seawater-based medium used for culturing these strains was prepared as follows. Coastal seawater was filtered using a 0.2-μm pore-size filter and autoclaved for 90 min. After autoclaving, seawater was sparged with 0.1 μm-filtered CO2 for 6 h followed by aeration overnight. After autoclaving and sparging, the seawater was amended with 1 mM NH_4_Cl, 1 μM FeCl_3_, mixed carbon sources (Cho and Giovannoni, 2014) and vitamins (Carini et al., 2013). Cultures were grown in one of the following conditions: (i) phosphate-addition (100 μM K_2_HPO_4_ added); or (ii) without additional phosphate (no phosphate added) (Zhang et al., 2019). Cultures were grown in the dark at 23° without shaking. The cells were not washed after pre-culture; instead, the bacteria were inoculated into the same medium (0.2% v/v) and the cell density was monitored using a Guava EasyCyte flow cytometer before harvesting (Merck Millipore, Billerica, MA, USA). Cell pellets were harvested from 10 ml culture by centrifugation which were stored at −80°C prior to lipid extraction.

### Lipid extraction and HPLC-MS

Lipid extraction from 10 ml bacterial culture was carried out using a modified Folch extraction method (Folch et al., 1957) using HPLC-grade chloroform (1 ml), Milli-Q water (0.3 ml) and LC-MS grade methanol (0.5 ml) in a 2 ml glass Chromacol vial (Smith et al., 2019). After phase separation by centrifugation, the lower chloroform phase containing the lipids was dried under nitrogen before resuspending in 0.5-1 mL of solvent (0.05 ml of 10 mM ammonium acetate in water, pH 9.2 and 0.95 ml acetonitrile). The lipid d17:1/12:0 sphingosylphosphoethanolamine (Sigma-Aldrich, 50 nM) was added to the samples and used as internal standard. Five μl of the lipid extract was injected onto the LC-MS and separated by a Dionex 3400RS HPLC using a hydrophilic interaction column (XBridge BEH amide XP column 2.5 μm 3.0×150 mm, Waters) according to their polar headgroup. Samples were run on a 15 min gradient from 95% (v/v) acetonitrile/5% (w/v) ammonium acetate (in water, 10 mM, pH 9.2) to 70% (v/v) acetonitrile/30% (w/v) ammonium acetate (in water, 10 mM, pH 9.2), followed by 5 minutes of isocratic run 70% acetonitrile/30% ammonium acetate. Ten minutes equilibration at the initial run condition were performed between samples. The flow rate was maintained at 150 μL min^-1^ and the column temperature at 30 °C. The injection volume was 5 μL for each run; the ionisation was done in both positive and negative mode. Drying conditions were the same for both modes (8 L min^-1^ drying gas at 300 °C; nebulising gas pressure of 15 psi), while the end cap voltage was 4,500 V in positive mode and 3,500 V in negative mode, both with 500 V offset. Identification of lipid classes was made through MS^n^ fragmentation using the amaZon SL ion trap mass spectrometer. Data analyses were carried out using the Bruker Compass Software with DataAnalysis for peak identification and lipid MS^n^ fragmentation and QuantAnalysis for lipid quantification against internal standard sphingosylphosphoethanolamine (SPE). The abundance of each lipid was normalized against sphingosylphosphoethanolamine and expressed as relative abundance against phosphatidylglycerol (PG).

### Bioinformatics and statistics

The genome sequences of strains RCA23 and LE17 and the single-cell amplified genomes (SAGs) of AB-661-I11, AB-661-L17 and AB-661-M21 were analysed using the JGI IMG portal (https://img.jgi.doe.gov/). Genomes were searched for the presence of the following genes involved in lipid renovation: *plcP*(MED193_17359), *olsA* (SPO1979), *glsB* (SPO2489) and *olsB* (SPO1980). Student*-t* tests were performed using Excel version 2017.

## Acknowledgements

This project has received funding from the European Research Council (ERC) under the European Union’s Horizon 2020 research and innovation programme (grant agreement no. 726116) and Deutsche Forschungsgemeinschaft within the Collaborative Research Centre Rosoebacter (TRR51).

